# A unified picture of neuronal avalanches arises from the understanding of sampling effects

**DOI:** 10.1101/759613

**Authors:** J. P. Neto, F. P. Spitzner, V. Priesemann

## Abstract

To date, it is still impossible to sample the entire mammalian brain with single-neuron precision. This forces one to either use spikes (focusing on few neurons) or to use coarse-sampled activity (averaging over many neurons, e.g. LFP). Naturally, the sampling technique impacts inference about collective properties. Here, we emulate both sampling techniques on a spiking model to quantify how they alter observed correlations and signatures of criticality. We discover a general effect: when the inter-electrode distance is small, electrodes sample overlapping regions in space, which increases the correlation between the signals. For coarse-sampled activity, this can produce power-law distributions even for non-critical systems. In contrast, spike recordings enable one to distinguish the underlying dynamics. This explains why coarse measures and spikes have produced contradicting results in the past – that are now all consistent with a slightly subcritical regime.

## 1 Introduction

For more than two decades, it has been argued that the cortex might operate at a critical point [1–6]. The criticality hypothesis states that by operating at a critical point, neuronal networks could benefit from optimal information-processing properties. Properties maximized at criticality include the correlation length [7], the autocorrelation time [6], the dynamic range [8, 9] and the richness of spatio-temporal patterns [10, 11].

Evidence for criticality in the brain often derives from measurements of **neuronal avalanches**. Neuronal avalanches are cascades of neuronal activity that spread in space and time. If a system is critical, the probability distribution of avalanche size *p*(*S*) follows a power law *p*(*S*) ∼ *S*^−*α*^[7, 12]. Such power-law distributions have been observed repeatedly in experiments since they were first reported by Beggs & Plenz in 2003 [1].

However, not all experiments have produced power laws and the criticality hypothesis remains controversial. It turns out that results for cortical recordings *in vivo* differ systematically:

Studies that use what we here call **coarse-sampled** activity typically produce power-law distributions [1, 13–22]. In contrast, studies that use **sub-sampled** activity typically do not [15, 23–27]. Coarse-sampled activity include LFP, M/EEG, fMRI and potentially calcium imaging, while sub-sampled activity is front-most spike recordings. We hypothesize that the apparent contradiction between coarse-sampled (LFP-like) data and sub-sampled (spike) data can be explained by the differences in the recording and analysis procedures.

In general, the analysis of neuronal avalanches is not straightforward. In order to obtain avalanches, one needs to define discrete events. While spikes are discrete events by nature, a coarse-sampled signal has to be converted into a binary form. This conversion hinges on thresholding the signal, which can be problematic [28–31]. Furthermore, events have to be grouped into avalanches, and this grouping is typically not unique [23]. As a result, avalanche-size distributions depend on the choice of the threshold and temporal binning [1, 32].

In this work, we show how thresholding and temporal binning interact with a (so far ignored) effect. Under coarse-sampling, neighboring electrodes may share the same field-of-view. This creates a distance-dependent *measurement overlap* so that the activity that is recorded at different electrodes may show *spurious correlations*, even if the underlying spiking activity is fully uncorrelated. We show that the inter-electrode distance may therefore impact avalanche-size distributions more severely than the underlying neuronal activity.

In the following, we explore the role of the recording and analysis procedures on a generic, locally-connected network of spiking neurons. We compare apparent signs of criticality under sub-sampling versus coarse-sampling. To that end, we vary the distance to criticality of the underlying system over a wide range, from uncorrelated (Poisson) to highly-correlated (critical) dynamics. We then derive signatures of criticality – as is done in experiments – and study how results depend on electrode distance and temporal binning.

## 2 Results

The aim of this study is to understand **how the sampling of neural activity** affects the inference of the underlying collective dynamics. It is not about introducing a novel model that might generate critical dynamics. Therefore, we use an established phenomenological model, where the distance to criticality can be precisely tuned. To study sampling effects, we use a two-level setup inspired by [35]: An underlying network model, on which activity is then *sampled* with a grid of 8 × 8 virtual electrodes. All parameters of the model, the sampling and the analysis are closely matched to those known from experiments (see Methods).

In order to evaluate sampling effects, we want to *precisely* set the underlying dynamics. Therefore, we employ the established branching model, which is well understood analytically [10, 26, 33–35]. Inspired by biological neuronal networks, we simulate the branching dynamics on a dense 2D topology with *N*_N_ = 160 000 neurons where each neuron is connected to *K* ≈ 1000 local neighbors. To emphasize the locality, the synaptic strength of connections decays with the distance *d*_N_ between neurons. For a detailed comparison with different topologies, see the Supplemental Information (Fig. S1).

### 2.1 The branching parameter *m* sets the distance to criticality

In order to compare apparent signatures of criticality with the true, underlying dynamics, we first give some intuition about the branching model. The **branching parameter** *m* quantifies the probability of *postsynaptic activations*, or in other words, how many subsequent spikes are caused (on average) by a single spike. With increasing *m* → 1, a single spike triggers increasingly long cascades of activity. These cascades determine the timescale over which fluctuations occur in the population activity – this **intrinsic timescale** *τ* describes the dynamic state of the system and its distance to criticality.

The intrinsic timescale can be analytically related to the branching parameter by *τ* ∼ −1/ln (*m*). As *m* → 1, *τ* → ∞ and the population activity becomes “bursty”. We illustrate this in Fig. 1B and Table 1: For Poisson-like dynamics (*m* ≈ 0), the intrinsic timescale is zero 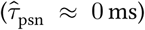 and the activity between neurons is uncorrelated. As the distance to criticality becomes smaller (*m* → 1), the intrinsic timescale becomes larger 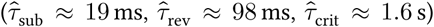, fluctuations become stronger, and the spiking activity becomes more and more correlated in space and time.

**Table 1:** Parameters and intrinsic timescales of dynamic states. All combinations of branching parameter *m* and per-neuron drive *h* result in a stationary activity of 1 Hz per neuron. Due to the recurrent topology, it is more appropriate to consider the measured autocorrelation time 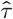 rather than the analytic timescale *τ*.

| State name | $m$ | $\hat{\tau}$ (measured) | $\tau = \frac{-2\text{ms}}{\ln m}$ | $h$ |
| --- | --- | --- | --- | --- |
| Poisson | 0.0 | $0.1 \pm 0.1$ ms | 0.0 ms | $2 \times 10^{-3}$ |
| Subcritical | 0.9 | $18.96 \pm 0.09$ ms | 18.9 ms | $2 \times 10^{-4}$ |
| Reverberating | 0.98 | $98.3 \pm 1.0$ ms | 98.9 ms | $4 \times 10^{-5}$ |
| Critical | 0.999 | $1.58 \pm 0.12$ s | 1.99 s | $2 \times 10^{-6}$ |

### 2.2 Avalanches are extracted differently under coarse-sampling and sub-sampling

At each electrode, we sample both the spiking activity of the closest neuron (sub-sampling) and a spatially averaged signal that emulates LFP-like coarse-sampling.

Both **sub-sampling** and **coarse-sampling** are sketched in Fig. 1A: For coarse-sampling (left), the signal from each electrode channel is composed of varying contributions (orange circles) of all surrounding neurons. The contribution of a particular spike from neuron *i* to electrode *k* decays as 1/*d*_*ik*_ with the neuron-to-electrode distance *d*_*ik*_ (see Supplemental Information for an extended discussion on the impact of the distance dependence). In contrast, if spike detection is applied (Fig. 1A, right), each electrode signal captures the spiking activity of few individual neurons (highlighted circles).

To test both recording types for criticality, we apply the standard analysis that provides a probability distribution *p*(*S*) of the *avalanche size S*: In theory, an avalanche describes a cascade of activity where individual units – here neurons – are consecutively and causally activated. Each activation is called an event. The avalanche size is then the total number of events in the time between the first and the last activation. A power law in the size distribution of these avalanches is a hallmark of criticality [6]. In practice, the actual size of an avalanche is hard to determine because individual avalanches are not clearly separated in time; the coarse-sampled signal is continuous-valued and describes the local population. In order to extract binary events for the avalanche analysis (Fig. 2), the signal has to be thresholded – which is not necessary for spike recordings, where binary events are inherently present as timestamps.

**Figure 1:**
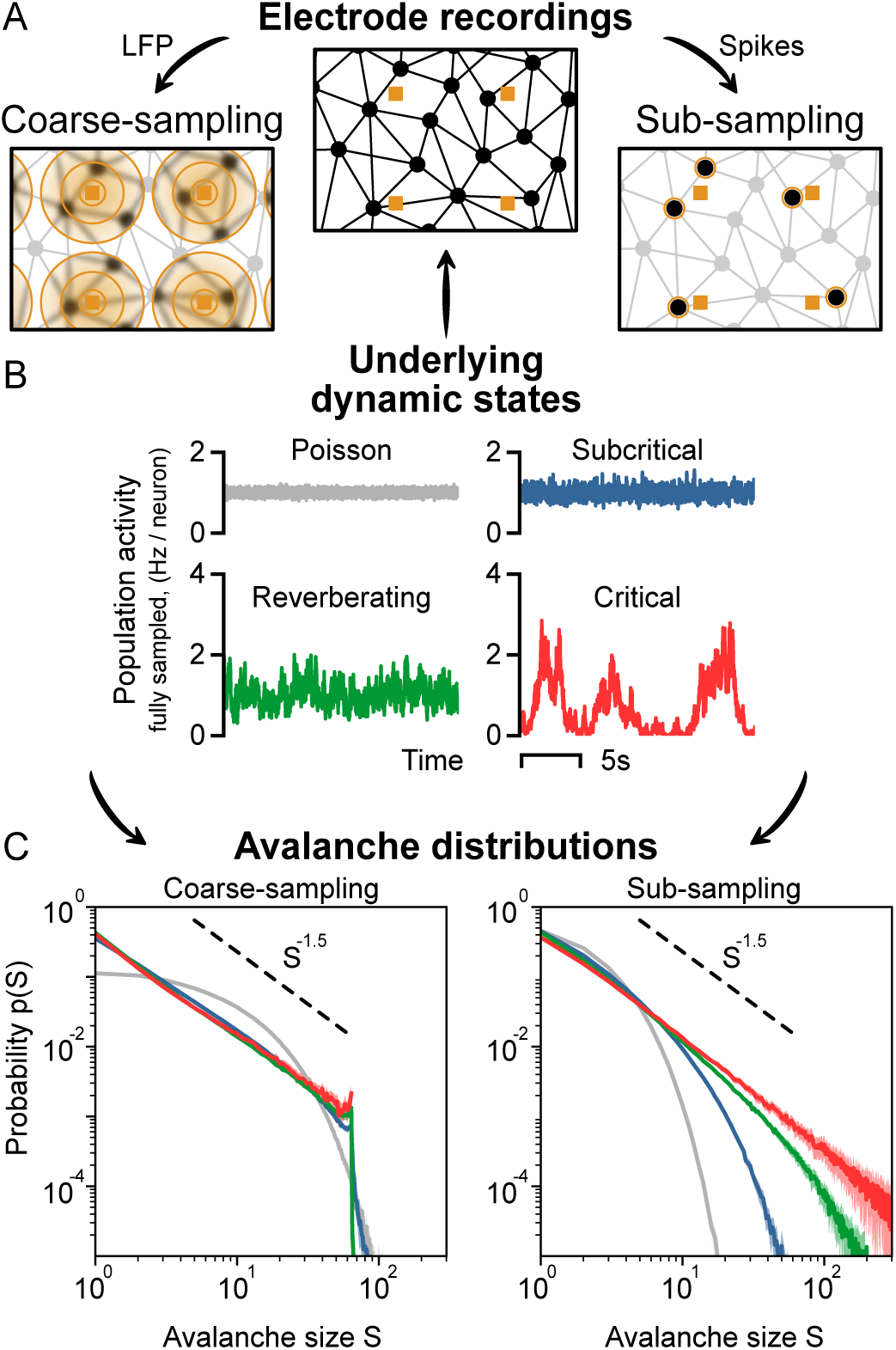
Sampling affects the assessment of dynamic states from neuronal avalanches. **A:** Representation of the sampling process of neurons (black circles) using electrodes (orange squares). Under coarse-sampling (e.g. LFP), activity is measured as a weighted average in the electrode’s vicinity. Under sub-sampling (spikes), activity is measured from few individual neurons. **B:** Fully sampled population activity of the neuronal network, for states with varying intrinsic timescales *τ*: Poisson 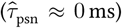, subcritical 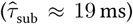, rever-berating 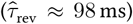 and critical 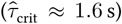. **C:** Avalanche-size distribution *p*(*S*) for coarse-sampled (left) and sub-sampled (right) activity. Sub-sampling allows for separating the different states, while coarse-sampling leads to *p*(*S*) ∼ *S*^−*α*^ for all states except Poisson. **Parameters**: Inter-electrode distance *d* _E_ = 400 µm and time-bin size Δ*t* = 8 ms.

**Figure 2:**
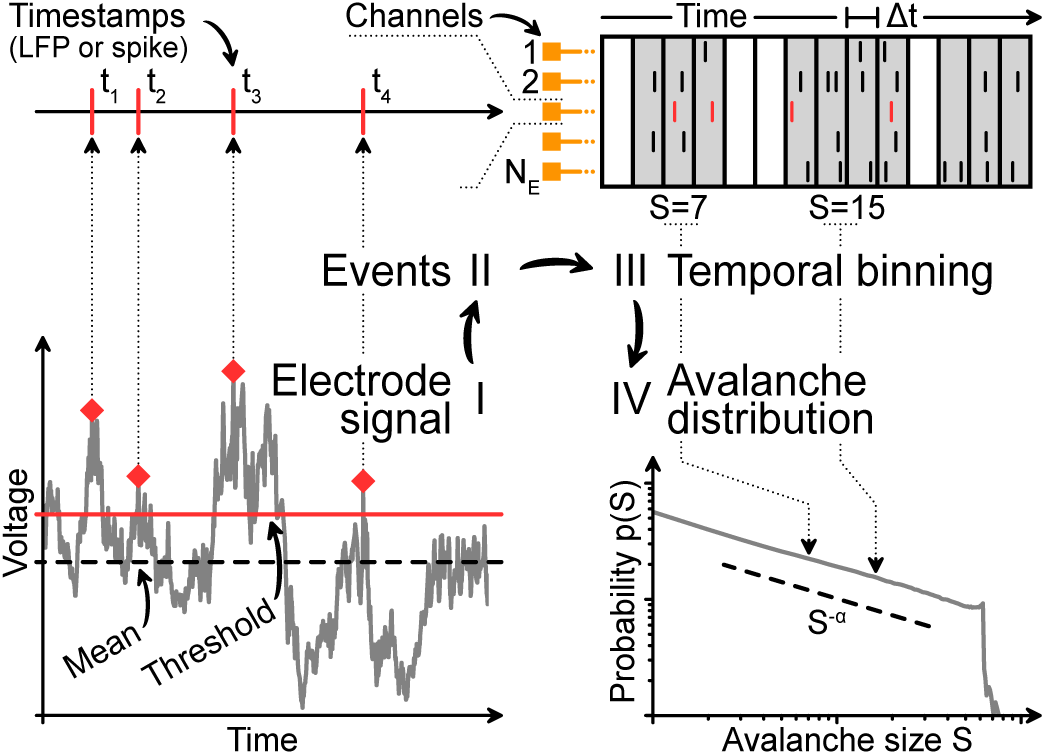
Analysis pipeline for avalanches from sampled data. **I:** Under coarse-sampling (LFP-like), the recording is demeaned and thresholded. **II:** The timestamps of events are extracted. Under sub-sampling(spikes), timestamps are obtained directly. **III:** Events from all channels are binned with time-bin size *Δt* and summed. The size *S* of each neuronal avalanche is calculated. **IV:** The probability of an avalanche size is given by the (normalized) count of its occurrences throughout the recording.

### 2.3 Coarse-sampling makes dynamic states indistinguishable

Irrespective of the applied sampling, the inferred avalanche distribution *should* represent the true dynamic state of the system.

However, under coarse-sampling (Fig. 1C, left), the avalanche-size distributions of the subcritical, reverberating and critical state are virtually indistinguishable. Intriguingly, all three show a power law. The observed exponent *α* = 1.5 is associated with a critical branching process. Only the uncorrelated (Poisson-like) dynamics produce a non-power-law decay of the avalanche-size distribution.

Under sub-sampling (Fig. 1C, right), each dynamic state produces a unique avalanche-size distribution. Only the critical state, with the longest intrinsic timescale, produces the characteristic power law. Even the close-to-critical, reverberating regime is clearly distinguishable and features a “subcritical decay” of *p*(*S*).

### 2.4 Measurement overlap causes spurious correlations

Why are the avalanche-size distributions of different dynamic states hard to distinguish under coarse-sampling? The answer is hidden within the cascade of steps involved in the recording and analysis procedure. Here, we separate the impact of the involved processing steps. Most importantly, we discuss the consequences of *measurement overlap* – which we identify as a key explanation for the ambiguity of the distributions under coarse-sampling.

In order to obtain discrete events from the continuous time series for the avalanche analysis, each electrode signal is filtered and thresholded, binned with a chosen time-bin size *Δt* and, subsequently, the events from all channels are stacked. This procedure is problematic because **(i)** electrode proximity adds spatial correlations, **(ii)** temporal binning adds temporal correlations, and **(iii)** thresholding adds various types of bias [28–30].

As a result of the involved analysis of coarse-sampled data, spurious correlations are introduced that are not present in sub-sampled data. We showcase this effect in Fig. 3, where the Pearson correlation coefficient between two virtual electrodes is compared for both the (thresholded and binned) coarse-sampled and sub-sampled activity. For the same parameters and dynamic state, coarse-sampling leads to larger correlations than sub-sampling.

Depending on the distance between electrodes, multiple electrodes might record activity from the same neuron. This **measurement overlap** (or volume conduction effect) increases the spatial correlations between electrodes – and because from the signals from multiple electrode channels are combined in the analysis, correlations can originate from measurement over-lap alone.

**Figure 3:**
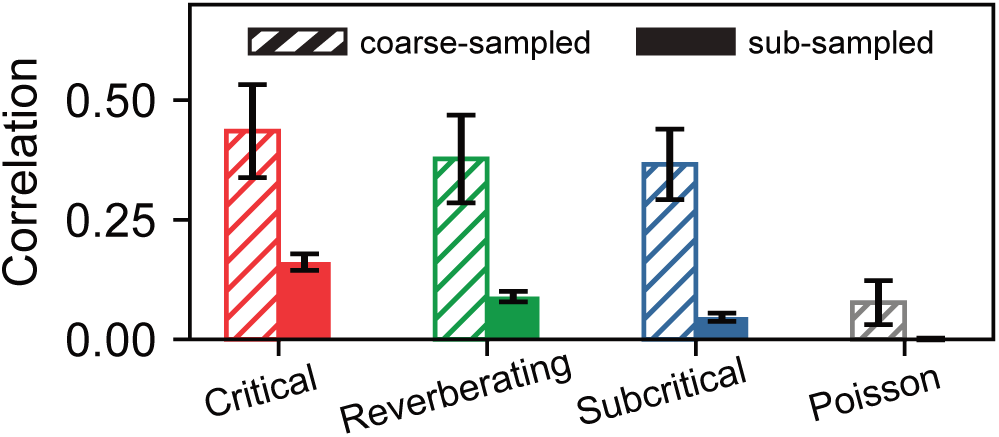
Coarse-sampling leads to greater correlations than sub-sampling. Pearson correlation coefficient between the signals of two adjacent electrodes for the different dynamic states. Even for independent (uncorrelated) Poisson activity, measured correlations under coarse-sampling are non-zero. **Parameters:** Inter-electrode distance *d*_E_ = 400 µm and time-bin size *Δt* = 8 ms.

### 2.5 Inter-electrode distance shapes criticality

Due to the measurement overlap, avalanche-size distributions under coarse-sampling depend on the inter-electrode distance *d*_E_ (Fig. 4A). For small inter-electrode distances, the overlap is strong. Thus, the spatial correlations are strong. Strong correlations manifest themselves in *larger* avalanches. However, under coarse-sampling the maximal observed size *S* of an avalanche is limited by the number of electrodes *N*_E_ [35]. This limit due to *N*_E_ manifests as a sharp cut-off and – in combination with spurious measurement correlations due to *d*_E_ – can shape the probability distribution. In the following, we show that these factors can be more dominant than the actual underlying dynamics.

In theory, supercritical dynamics are characterized by a *sharp peak* in the avalanche distribution at *S* = *N*_E_. Independent of the underlying dynamics, such a peak can originate from small electrode distances (Fig. 4A, *d*_E_ = 100 µm):

Avalanches are likely to span the small area covered by the electrode array. Furthermore, due to strong measurement overlap, individual events of the avalanche may contribute strongly to multiple electrodes.

Subcritical dynamics are characterized by a *pronounced decay* already for *S* < *N*_E_. Independent of the underlying dynamics, such a decay can originate from large electrode distances (Fig. 4A, *d*_E_ = µm): Locally propagating avalanches are unlikely to span the large area covered by the electrode array. Furthermore, due to the weaker measurement overlap, individual events of the avalanche may contribute strongly to one electrode (or to multiple electrodes but only weakly).

Consequently, there exists a *sweet-spot* value of the interelectrode distance *d*_E_ for which *p*(*S*) appears convincingly critical (Fig. 4A, *d*_E_ = 250 µm): a power law *p*(*S*) ∼ *S*^−*α*^ spans all sizes up to the cut-off at *S* = *N*_E_. However, the dependence on the underlying dynamic state is minimal.

Independently of the apparent dynamics, we observe the discussed cut-off at *S* = *N*_E_, which is also often seen in experiments (Fig. 5). Note, however, that this cut-off only occurs under coarse-sampling (see again Fig. 1C). When spikes are used instead (Fig. 6), the same avalanche can reach an electrode repeatedly in quick succession – whereas such double-events are circumvented when thresholding at the population level. For more details see Fig. S2.

A further signature of criticality is obtained by estimating the branching parameter. This is traditionally done at the avalanche level: The *estimated branching parameter* of the neuronal avalanches, *m*_av_, is defined as the average ratio of events between subsequent time bins in an avalanche, i.e. during non-zero activity [1, 32]. Note that, due to coalescence and drive effects, *m*_av_ can differ from *m* proper [23, 34].

Obtaining *m*_av_ for different electrode distances results in a picture consistent with the one from avalanche-size distributions (Fig. 4B). In general, the dependence on the electrode distance is stronger than the dependence on the underlying state. At the particular value of the inter-electrode distance where *m*_av_ = 1, the distributions appear critical. If *m*_av_ < 1 (*m*_av_ > 1), the distributions appear subcritical (supercritical). Because the probability distributions and the estimated branching parameter share this dependence, a wide range of dynamic states would be consistently misclassified – solely as a function of the inter-electrode distance.

**Figure 4:**
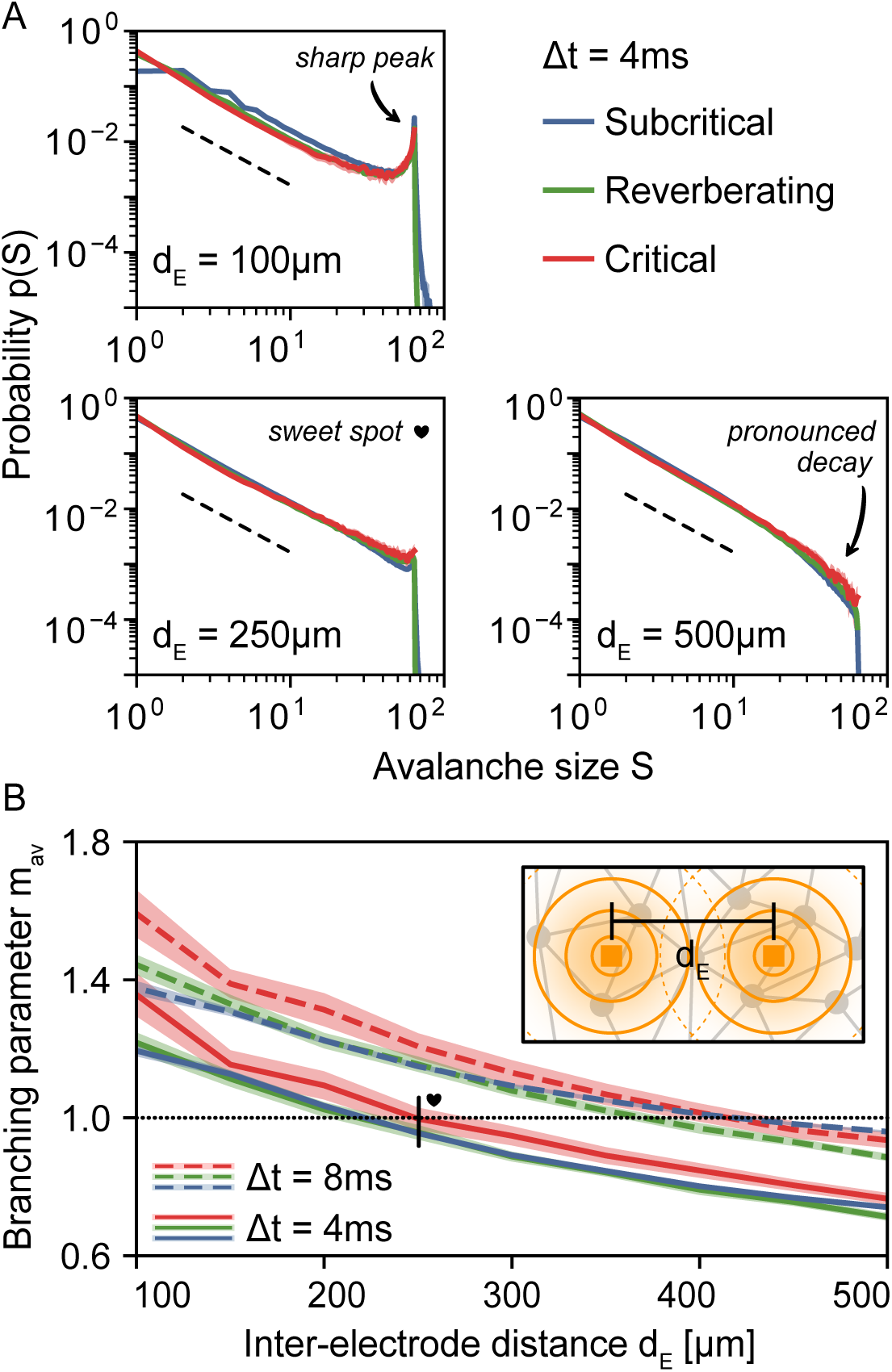
Under coarse-sampling, apparent dynamics depend on the inter-electrode distance *d*_E_. **A**: For small distances (*d*_E_ = 100 μm), the avalanche-size distribution *p*(*S*) indicates (apparent) supercritical dynamics: *p*(*S*) ∼ *S*^−*α*^ with a *sharp peak* near the electrode number *N*_E_ = 64. For large distances (*d*_E_ = 500 μm), *p*(*S*) indicates subcritical dynamics: *p*(*S*) ∼ *S*^−*α*^ with a pronounced decay already for *S* < *N*_E_. There exists a *sweet-spot* value (*d*_E_ =250 µm) for which *p*(*S*) indicates critical dynamics: *p*(*S*) ∼ *S*^−*α*^ until the the cut-off is reached at *s=N*_E_. The particular sweet-spot value of *d*_E_ depends on time-bin size (here, *Δt* = 4 ms). As a guide to the eye, dashed lines indicate *S*^-1.5^ B: The branching parameter *m*_av_ is also biased by *d* when estimated from neuronal avalanches. Apparent criticality (*m*_av_ ≈ 1, dotted line) is obtained with *d*_E_ = 250 µm and *Δt* = 4 ms but also with *d*_E_ = 400 µm and *Δt* = 8 ms. **B, Inset:** representation of the measurement overlap between neighboring electrodes; when electrodes are placed close to each other, spurious correlations are introduced.

### 2.6 Temporal binning determines scaling exponents

Apart from the inter-electrode distance, the choice of temporal discretization that underlies the analysis may alter avalanche-size distributions. This *time-bin size Δt* varies from study to study and it can severely impact the observed distributions [1, 23, 36, 37]. With smaller bin sizes, avalanches tend to be separated into small clusters, whereas larger bin sizes tend to “glue” subsequent avalanches together [23]. Interestingly, this not only leads to larger avalanches, but specifically to *p*(*S*) ∼ *S*^−*α*^, where the exponent *α* increases systematically with bin size [1, 36]. Such a changing exponent is not expected for conventional systems that self-organize to criticality: Avalanches would be *separated in time*, and *α* should be fairly bin-size invariant for a large range of *Δt* [23, 37, 38].

Our coarse-sampled model reproduces these characteristic experimental results (Fig. 5). It also reproduces the previously reported scaling [1] of the exponent with bin size *α* ∼ *Δt*^−*β*^ (Fig. 5, insets). Except for the Poisson dynamics, all the model distributions show power laws. Moreover the distributions are strikingly similar, not just to the experimental results, but also to each other. This emphasizes how sensitive signs of criticality are to analysis parameters: All the shown dynamic states are consistent with the ubiquitous avalanche-size distributions that are observed in coarse-sampled experiments.

When spikes are used instead, power-law distributions only arise from critical dynamics. For comparison with the coarse-sampled results in Fig. 5, we show avalanche-size distributions from experimental spike recordings and sub-sampled simulations in Fig. 6. In this case, power laws are produced only by in vitro cultures and the simulations that are (close-to) critical. In vivo spike recordings on awake subjects and simulations of subcritical dynamics produce distributions that feature a pronounced decay instead of power laws. In contrast to coarse-sampling, the avalanche distributions that stem from sub-sampled measures (spikes) allow us to clearly tell apart the underlying dynamic states from one another.

Overall, as our results on coarse-sampling have shown, different sources of bias – here the measurement overlap and the bin size – can perfectly outweigh each other. For instance, smaller electrode distances (that increase correlations) can be compensated by making the time-bin size smaller (which again decreases correlations). This was particularly evident in Fig. 4B, where increasing *d*_E_ could be outweighed by increasing *Δt* in order to obtain a particular value for the branching parameter *m*_av_. The same relationship was again visible in Fig. 5C-F: For the shown *d*_E_ = 400 µm (see also Fig. S6 for *d*_E_ = 200 µm), only *Δt* = 8 ms results in *α* = 1.5 – the correct exponent for the underlying dynamics. Since the electrode distance cannot be varied in most experiments, selecting anything but the one “lucky” *Δt* will cause a bias.

**Figure 5:**
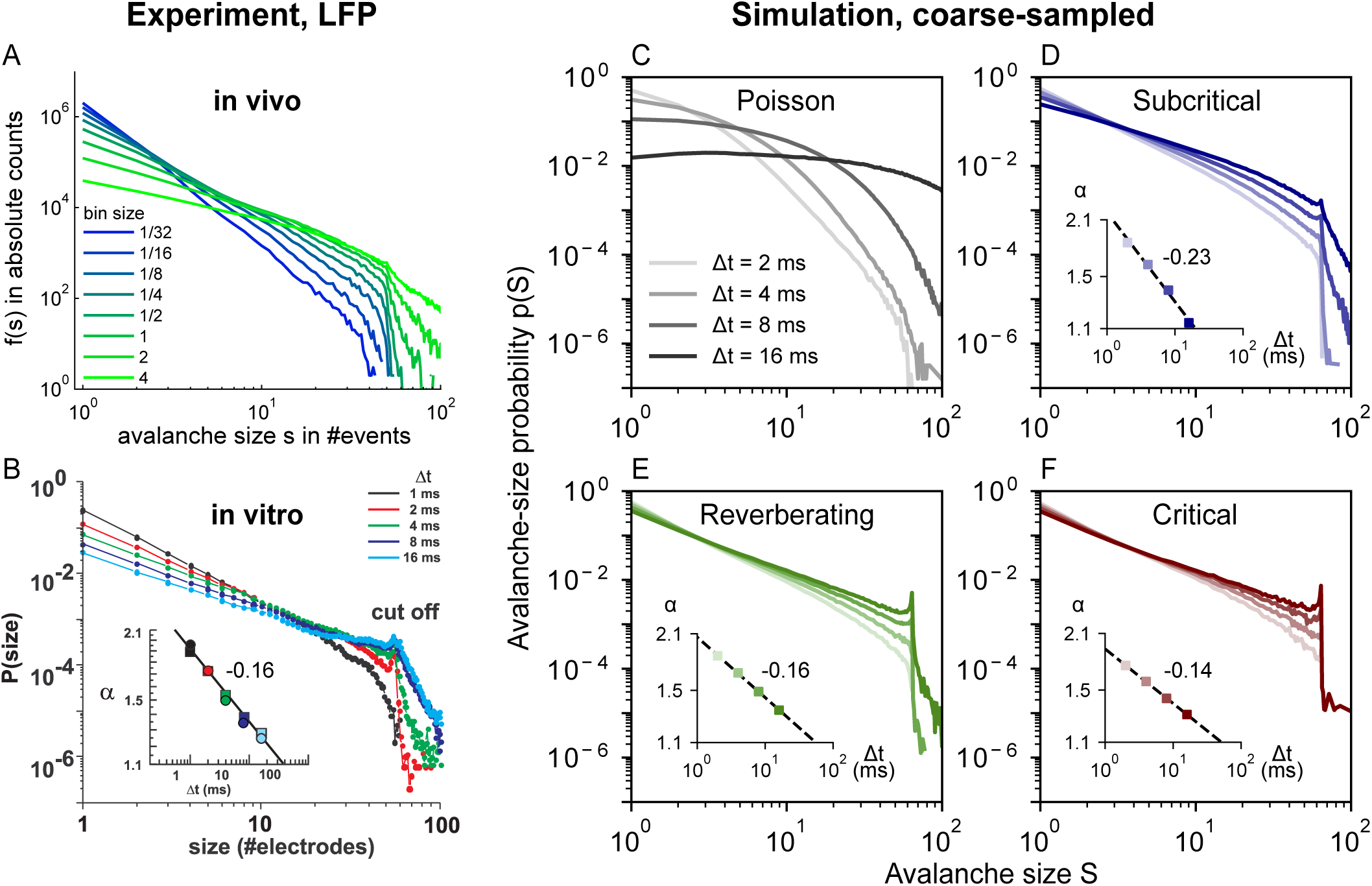
In vivo and in vitro avalanche-size distributions *p*(*S*) from LFP depend on time-bin size *Δt*. Experimental LFP results are reproduced by many dynamics states of coarse-sampled simulations. **A**: Experimental *in vivo* results (LFP, human) from an array of 60 electrodes, adapted from [36]. **B:** Experimental *in vitro* results (LFP, culture) from an array with 60 electrodes, adapted from [1]. **C–F**: Simulation results from an array of 64 virtual electrodes and varying dynamic states, with time-bin sizes between 2 ms ≤ *Δt* ≤ 16 ms and *d*_E_ = 400 µm. Subcritical, reverberating and critical dynamics produce power-law distributions with bin-size-dependent exponents *α*. **Insets:** Distributions are fitted to *p*(*S*) ∼ *S*^−*α*^. The magnitude of *α* decreases as *Δt*^−*β*^ with −*β* indicated next to the insets.

### 2.7 Scaling laws fail under coarse-sampling

The most used indication of criticality in neuronal dynamics is the avalanche-size distribution *p*(*S*). However, at criticality, the *avalanche duration distribution p*(*D*) and the *average avalanche size* for a given duration, ⟨*S*⟩(*D*), should also follow power-laws, each with a respective *critical exponent* [12]:

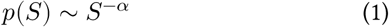

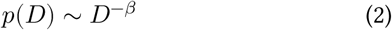

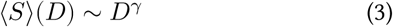

The exponents are related to one another by the scaling relationship

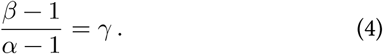

For a pure branching process – or any process in the mean-field directed percolation universality class [7, 39] – they take the values *α* = 3/2, *β* = 2 and *γ* = 2.

Lastly, at criticality, avalanches of vastly different duration still have the same *average shape*: The activity *s*(*t, D*) at any given time *t* (within the avalanche’s lifetime *D*) is described by a universal scaling function *ℱ*, so that

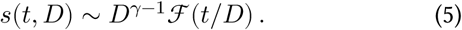

In other words, changing *s*(*t, D*) → *s*(*t, D*)/*D*^*γ*−1^ and *t* → *t*/*D* should result in a data collapse for the average avalanche shapes of all durations.

From Eqs. (3)–(5), we have three independent ways to determine the exponent *γ*. Consistency between the three is a further test of criticality. However, to the best of our knowledge, experimental evidence with the full set of scaling laws was only observed under sub-sampling: from spikes of in vitro recordings [40, 41].

The absence of scaling laws in coarse-sampled data can be explained by how coarse-sampling biases the average shape: the cut-off in *p*(*S*) near the number of electrodes *S* = *N*_E_ implies that ⟨*S*⟩(*D*) < *N*_E_. From Eq. (3) we have 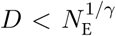. If *γ* > 1 the cut-off in *p*(*S*) causes a much earlier cut-off in both *p*(*D*) and ⟨*S*⟩(*D*).

Given that experiments typically have *N*_E_ ∼ 10^2^ electrodes, *p*(*D*) of a pure branching process (with) would span a power-law for less than one order of magnitude. However, the typical standard to reliably fit a power-law is at least two orders of magnitude [42]. While this is problematic under coarse-sampling (Fig. 5), we have shown that the hard cut-off is not present under sub-sampling (Fig. 6).

Again comparing the two ways of sampling, we now apply the independent measurements of *γ* to our model with critical dynamics (Fig. 7). We find consistent exponents under sub-sampling.

In this case, although they differ from those expected for a pure branching process (*γ* = 2), the exponents we find are compatible with the experimental values of *γ*_exp_ = 1.3 ± 0.05 reported in [40] and 1.3 ≤ *γ*_exp_ ≤ 1.5 reported in [41].

Under coarse-sampling, however, the exponent obtained from the shape collapse (*γ* ≈ 0.74) greatly differs from the other two (*γ* ≈ 1.74, *γ* ≈ 1.62), Fig. 7F. Moreover, the extremely short range available to fit *p*(*D*) and ⟨*S*⟩(*D*) with power-laws (1 ≤ *D* ≤ 6) makes the estimated exponents unreliable.

To conclude, the full set of critical exponents revealed criticality only under sub-sampling. Only in this case we observed both, a match between all the measurements of the exponent *γ*, and a power-law behavior extending over a range large enough to reliably fit them.

**Figure 6:**
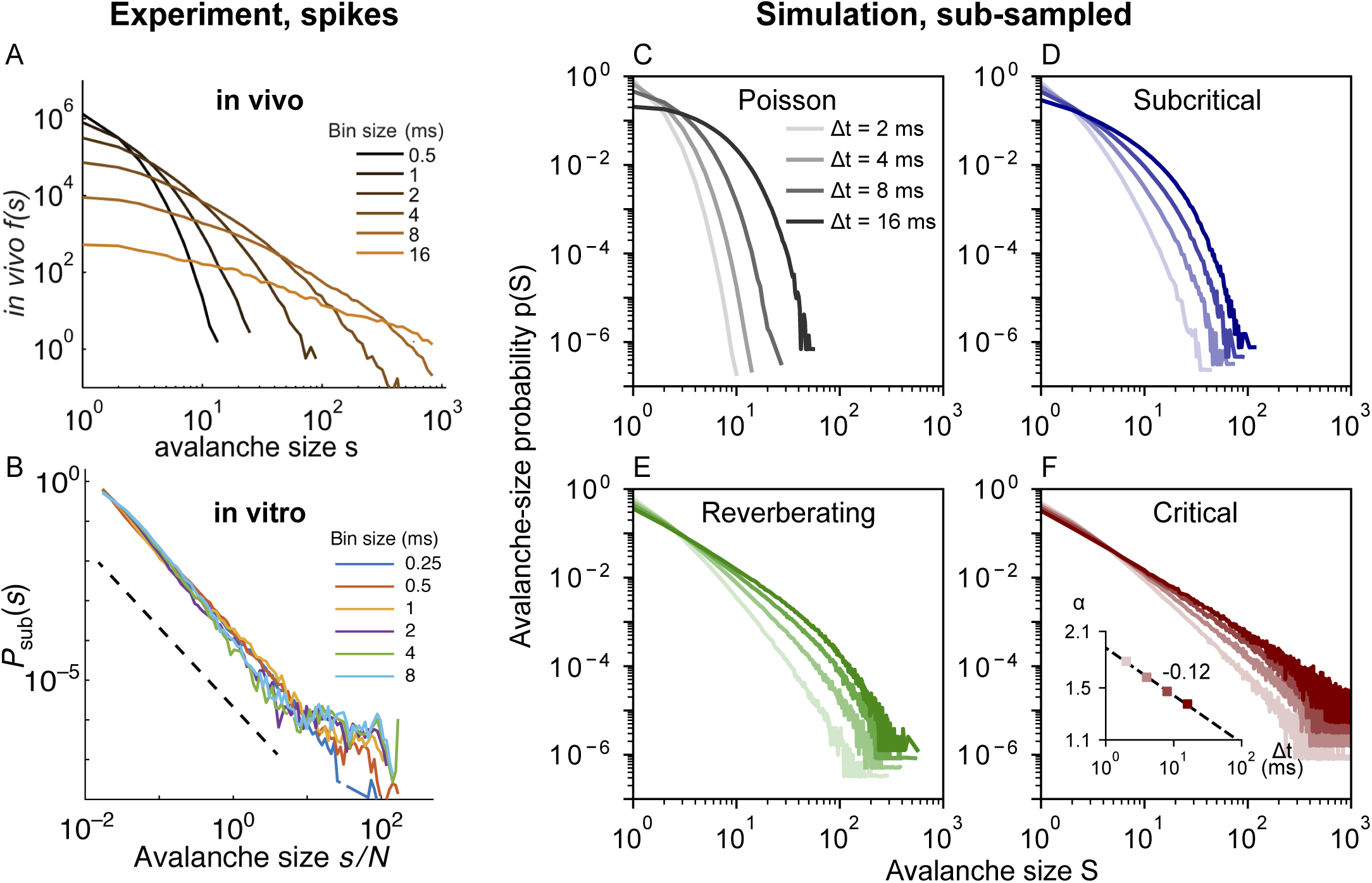
*In vivo* avalanche-size distributions *p*(*S*) from spikes depend on time-bin size *Δt*. *In vivo* results from spikes are reproduced by sub-sampled simulations of subcritical to reverberating dynamics. Neither spike experiments nor sub-sampled simulations show the cut-off that is characteristic under coarse-sampling. **A**: Experimental *in vivo* results (spikes, awake monkey) from an array of 16 electrodes, adapted from [23]. The pronounced decay and the dependence on bin size indicate subcritical dynamics. **B:** Experimental *in vitro* results (spikes, culture DIV 34) from an array with 59 electrodes, adapted from [37]. Avalanche-size distributions are independent of time-bin size and produce a power law over four orders of magnitude. In combination, this indicates critical dynamics with a separation of timescales. **C–F**: Simulation for sub-sampling, analogous to Fig. 5. Subcritical dynamics do not produce power-law distributions and are clearly distinguishable from critical dynamics. **F**: Only the (close-to) critical simulation produces power-law distributions. Note the dependence on time-bin size: In contrast to the *in vitro* culture, the simulation does not feature a separation of time scales (due to external drive and stationary activity) which causes a bin-size dependence.

**Figure 7:**
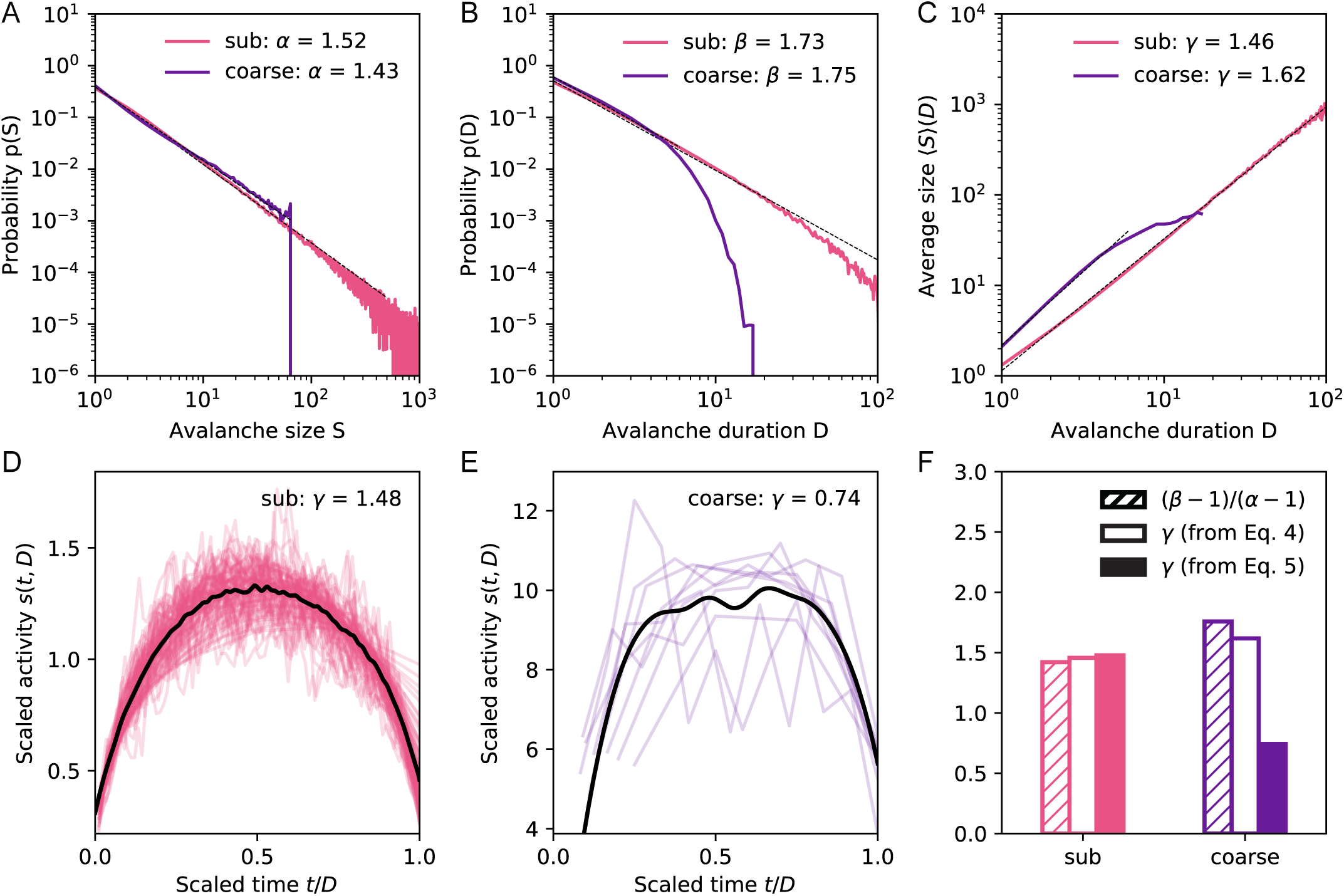
Scaling laws of a system with critical dynamics under coarse- and sub-sampling. **A–C**: Avalanche-size distribution *p*(*S*) ∼ *S*^−*α*^, avalanche-duration distribution *p*(*D*) ∼ *D*^−*β*^, and average size for a given duration ⟨*S*⟩(*D*) ∼ *D*^*γ*^, respectively, for sub-sampled (“sub”) and coarse-sampled (“coarse”) simulations. Distributions under sub-sampling easily span more than one order of magnitude, while coarse-sampled distributions suffer from an early cut-off (which hinders power-law fits). **D, E:** Shape collapse of *s*(*t, D*) ∼ *D*^*γ*−1^ *ℱ*(*t*/*D*) for sub-sampled and coarse-sampled data, respectively. Under coarse-sampling, the early duration cut-off results in few unique shapes for the collapse (corresponding to unique -values). **F**: Comparison of the critical exponents obtained independently from Eqs. (3)–(5). Exponents are consistent only under sub-sampling. **Parameters:** *d*_E_ = 400 µm and *Δt* = 8 ms.

## 3 Discussion

When inferring collective network dynamics from partially sampled systems, it is crucial to understand how the sampling biases the measured observables. Without this understanding, an elaborate analysis procedure – such as the one needed to study neuronal avalanches from coarse-sampled data – can result in a misclassification of the underlying dynamics.

We have shown that the analysis of neuronal avalanches based on (LFP-like) coarse-sampled data can produce indistinguishable results for systems with vastly different spatiotemporal signatures. These signatures derive from underlying dynamic states that, in this work, range from subcritical to critical – a range over which the intrinsic timescale undergoes a hundred-fold increase. And yet, the resulting avalanche-size distributions can be uninformative and ambiguous (Fig. 1).

The ambiguity of neuronal avalanches partially originates from spurious correlations. We have demonstrated the generation of spurious correlations from two sampling and processing mechanisms: measurement overlap (due to volume conduction) and temporal binning. Other studies found further mechanisms that can generate apparent power-law distributions by (purposely or accidentally) introducing correlations into the *observed* system. For instance, correlated input introduces temporal correlations already into the *underlying* system [43, 44]. Along with thresholding and low-pass frequency filtering – which add temporal correlations to the *observed* system [24, 45] – this creates a large space of variables that either depend on the system, sampling and processing, or a combination of both.

As our results focus on sampling and processing, we believe that the observed impact on avalanche-size distributions is general and model independent. We deliberately chose a simple model and confirmed that our results are robust to parameter changes: For instance, employing a more realistic topology causes no qualitative difference (Fig. S1). Furthermore, as a proof of concept, we checked the impact of measurement overlap in the 2D Ising model (Fig. S3). Even in such a fundamental model a measurement overlap can bias the assessment of criticality.

With our results on sampling effects, we can revisit the previous literature on neuronal avalanches. In the model, we found that coarse-sampling clouds the differences between subcritical, reverberating, and critical dynamics: The avalanche distributions always resemble power laws (Fig. 1). Because of this ambiguity, the power-law distributions obtained ubiquitously from LFP, EEG, MEG and BOLD activity should be taken as evidence of neuronal activity with spatio-temporal correlations – but not necessarily of criticality proper; the coarse-sampling hinders such a precise classification.

In contrast, a more precise classification is possible under sub-sampling. If power-law distributions are observed from (sub-sampled) spiking activity, they do point to critical dynamics. For spiking activity, we even have mathematical tools to infer the precise underlying state in a sub-sampling-invariant manner that does not rely on avalanche distributions [26, 46]. Having said so, not all spike recordings point to critical dynamics: While in vitro recordings typically do produce power-law distributions [37, 40, 47, 48], recordings from awake animals do not [15, 17, 23, 49]. Together, these results suggest that in vitro systems self-organize towards criticality, whereas the cortex of awake animals (and humans) operates *near criticality* – in a slightly subcritical, reverberating *regime*.

The reverberating regime harnesses benefits associated with criticality, and it unifies both types of in vivo results: For experiments on awake animals, spike-based studies indicate subcritical dynamics. While coarse measures produce power laws that indicate criticality, with this study we showed that they cannot distinguish critical from subcritical dynamics. Consistent with both, a brain that operates in a regime – as opposed to a fixed dynamic state – can flexibly tune response properties. In particular, the reverberating regime covers a specific range of dynamics in the vicinity of the critical point, where small changes in effective synaptic strength cause major changes in response properties. Hence, the reverberating regime is an ideal baseline [26] from which brain areas or neural circuits can adapt to meet task demands [36, 50–56].

In conclusion, our results methodically separate sampling effects from the underlying dynamic state. They overcome the discrepancy between the coarse-sampled and sub-sampled results of neuronal avalanches from awake animals. By offering a solution to a long-standing (critical) point of conflict, we hope to move beyond just describing a system as critical or not, and appreciate the richness of dynamic states around criticality.

## Acknowledgments

JPN, FPS and VP received financial support from the Max Planck Society. JPN received financial support from the Brazilian National Council for Scientific and Technological Development (CNPq) under Grant No. 206891/2014-8. We thank Jordi Soriano, and all members of our group, for valuable input. We thank Johannes Zierenberg and Bettina Royen for careful proofreading of the manuscript.

## Author contributions

J.P.N., F.P.S. and V.P. designed research; J.P.N., F.P.S. and V.P. performed research; J.P.N. and F.P.S. analyzed data; J.P.N., F.P.S. and V.P. wrote the paper.

## 4 Methods

### 4.1 Model Details

Our model is comprised of a two-level configuration, where a 2D network of *N*_N_ =160000 spiking neurons is sampled by a square array of *N*_E_ =8 × 8 virtual electrodes. Neurons are distributed randomly in space (with periodic boundary conditions) and, on average, nearest neighbors are *d*_N_ = 50 µm apart. While the model is inherently unit-less, it is more intuitive to assign some length scale – in our case the inter-neuron distance *d*_N_ – to set that scale: all other size-dependent quantities can then be expressed in terms of the chosen *d*_N_. For instance, the linear system size can be derived by realizing that the random placement of neurons corresponds to an ideal gas. It follows that 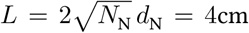 for uniformly distributed neurons. (For comparison, on a square lattice, the packing ratio would be higher and it is easy to see that the system size would be 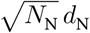.) Given the system size and neuron number, the overall neuronal density is *ρ* = 100/mm^2^. With our choice of parameters, the model matches typical experimental conditions in terms of inter-neuron distance, system size and neuron density (see Table 2 for details). The implementation of the model in C++, and the python code used to analyze the data and generate the figures, are available online at https://github.com/Priesemann-Group/criticalavalanches.

### 4.2 Topology

We consider a topology that enforces *local* spreading dynamics. Every neuron is connected to all of its neighbors within a threshold distance *d*_max_. The threshold is chosen so that on average *k* = 10^3^ outgoing connections are established per neuron. We thus seek the radius d_max_. of a disk whose area contains *k* neurous. Using the already known neuron density, we find 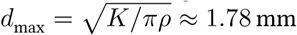. For every established connection, the probability of a recurrent activation decreases with increasing neuron distance. Depending on the particular distance *d*_*ij*_between the two neurons *i* and *j*, the connection has a normalized weight 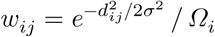 (with normalization constant 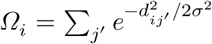. Our weight definition approximates the distance dependence of average synaptic strength. The parameter *σ* sets the *effective* distance over which connections can form (*d*_max_ is an upper limit for *σ* and mainly speeds up computation.) In the limit *σ* ⟶ ∞, the network is all-to-all connected. In the limit *σ* ⟶ 0, the network is completely disconnected. Therefore, the effective connection length *σ* enables us to fine tune *how local* the dynamic spreading of activity is. In our simulations, we choose *σ* = 6*d*_N_ = 300 μm. Thus, the overall reach is much shorter than *d*_max_ (*σ* ≈ 0.06 *d*_max_).

### 4.3 Dynamics

To model the dynamic spreading of activity, time is discretized to a chosen simulation time step, here *δt* = 2 ms, which is comparable to experimental evidence on synaptic transmission [57]. Our simulations run for 10^6^ time steps on an ensemble of 50 networks for each configuration (combination of parameters and dynamic state). This corresponds to ∼ 277 hours of recordings for each dynamic state.

The activity spreading is modeled using the dynamics of a branching process with external drive [26, 33]. At every time step *t*, each neuron *i* has a state *S*_*i*_(*t*) = 1 (spiking) or 0 (quiescent). If a neuron is spiking, it tries to activate its connected neighbors – so that they will spike in the next time step. All of these recurrent activations depend on the *branching parameter m*: Every attempted activation has a probability *p*_*ij*_ = *m w*_*ij*_ to succeed. (Note that the distance-dependent weights are normalized to 1 but the activation probabilities are normalized to *m*.) In addition to the possibility of being activated by its neighbors, each neuron has a probability *h* to spike spontaneously in the next time step. After spiking, a neuron is reset to quiescence in the next time step if it is not activated again.

Our model gives us full control over the dynamic state of the system – and its distance to criticality. The dynamic state is described by the *intrinsic timescale*. We can analytically calculate the intrinsic timescale τ =−*δt*/ ln (*m*), where *δt* is the duration of each simulated time step. Note that *m* – the control parameter that *tunes the system* – is set on the neuron level while *τ* is a (collective) network property (that in turn allows us to deduce an *effective m*). As the system is pushed more towards criticality (by setting *m* ⟶ 1), the intrinsic timescale diverges *τ* ⟶ ∞.

For consistency, we measure the intrinsic timescale during simulations. To that end, the (fully sampled) population activity at each time step is given by the number of active neurons *A*(*t*) = ∑_*i*_ *s*_*i*_ (*t*). A linear least-squares fit of the autoregressive relation 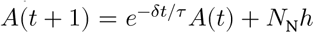 over the full simulated time series yields an estimate 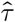 that describes each particular realization.

By adjusting the branching parameter *m* (setting the dynamic state) and the probability for spontaneous activations *h* (setting the drive), we control the distance to criticality *and* the average stationary activity. The activity is given by the *average spike rate r* = *h*/(*δt*(1−*m*)) of the network. For all simulations, we fix the rate to *r* = 1Hz Hz in order to avoid rate effects when comparing different states (see Table 1 for the list of parameter combinations). Note that, due to the non-zero drive *h* and the desired stationary activity, the model cannot be perfectly critical (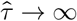, see Table 1).

### 4.4 Coalescence Compensation

With our probability-based update rules, it may happen that target neurons are simultaneously activated by multiple sources. This results in so-called *coalescence effects* that are particularly strong in our model due to the local activity spreading [34]. For instance, naively setting *m* = 1 (with *σ* = 300 µm)would result in an effective (measured) 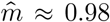, which has considerably different properties. Compared to e.g. *m* = 0.999 this would result in a 20-fold decrease in *τ.*

In order to compensate these coalescence effects, we apply a simple but effective fix: If an activation attempt is successful but the target neuron is already marked to spike in the next time step, another (quiescent) target is chosen. Because our implementation stores all the connected target neurons as a list sorted by their distance to the source, it is easy to activate the next neuron in that list. Thereby, the equivalent probability of the performed activation is as close to the originally attempted one as possible.

**Table 2:** Values and descriptions of the model parameters.

| Symbol | Value | Description |
| --- | --- | --- |
| $\Delta t$ | 2 – 16 ms | Time-bin size (duration) for temporal binning |
| $\Theta_k$ | 3 | Activity threshold, in units of standard deviations of the time series of electrode $k$ |
| $\delta t$ | 2 ms | Simulation time step |
| $r$ | 1 Hz | Average spike rate |
| $N_N$ | $1.6 \times 10^5$ | Number of neurons |
| $d_N$ | 50 $\mu\text{m}$ | Inter-neuron distance (measured between nearest neighbors) |
| $L$ | 4 cm | Linear system size |
| $\rho$ | 100/$\text{mm}^2$ | Neuronal density |
| $K$ | 1000 | Average network degree (outgoing connections per neuron) |
| $d_{\text{max}}$ | 1.78 mm | Connection length; all neurons within $d_{\text{max}}$ are connected |
| $\sigma$ | 300 $\mu\text{m}$ | Effective length of synaptic connections, sets the distance-dependence of the probabilities of recurrent activations |
| $N_E$ | $8 \times 8$ | Number of electrodes |
| $d_E$ | 50 – 500 $\mu\text{m}$ | Inter-electrode distance |
| $d_E^*$ | 10 $\mu\text{m}$ | Dead-zone around each electrode (no neurons present) |

### 4.5 Virtual Electrode Recordings

Our simulations are designed to mimic sampling effects of electrodes in experimental approaches. To simulate sampling, we use the readout of *N*_E_ = 64 virtual electrodes that are placed in 8×8 grid. Electrodes are separated by an inter-electrode distance that we specify in multiples of inter-neuron distance *d*_N_. It is kept constant for each simulation and we study the impact of the inter-electrode distance by repeated simulations spanning electrode distances between 1*d*_N_ = 50 µm and 10*d*_N_ = 500 µm. The electrodes are modeled to be point-like objects in space that have a small dead-zone of 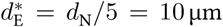 around their origin. Within the dead-zone, no signal can be recorded (in fact, we implement this by placing the electrodes first and the neurons second – and forbid neuron placements too close to electrodes.)

Using this setup, we can apply sampling that emulates either the detection of spike times or LFP-like recordings. To model the detection of spike times, each electrode only observes the single neuron that is closest to it. Whenever this particular neurons spikes, the timestamp of the spike is recorded. All other neurons are neglected – and the dominant sampling effect is *sub-sampling*. On the other hand, to model LFP-like recordings, each electrode integrates the spiking of all neurons *i* to electrode *k*,decays as 1/*d*_*ik*_ with the neuron-to-electrode distance. (Changing the dependence to 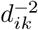 has no qualitative impact on the results.) The total signal of the electrode at time *t* is then 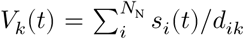 (Diverging electrode signals are prevented by the forbidden zone around the electrodes.) For such coarse-sampled activity, all neurons contribute to the signal and the contribution is weighted by their distance.

### 4.6 Avalanches

Taking into account all 64 electrodes, a new avalanche starts (by definition [1]) when there is at least one event (spike) in a time bin – given there was no event in the previous time bin (see Fig. 2). An avalanche ends whenever an empty bin is observed (no event over the duration of the time bin). Hence, an avalanche persists for as long as every consecutive time bin contains at least one event – which is called the *avalanche duration D* From here, it is easy to count the total number of events that were recorded across all electrodes and included time bins – which is called the *avalanche size*. The number of occurrences of each avalanche size (or duration) are sorted into a histogram that describes the avalanche distribution.

### 4.7 Analysis of Avalanches under Coarse and Sub-sampling

We analyze avalanche size distributions in a way that is as close to experimental practice as possible (see Fig. 2). From the simulations described above, we obtain two outputs from each electrode: a) a list containing spike times of the single closest neuron and b) a time series of the integrated signal to which all neurons contributed.

In case of the (sub-sampled) spike times a), the spiking events are already present in binary form. Thus, to define a neural avalanche, the only required parameter is the size of the time bin *Δt*(for instance, we may choose *Δt* = 4 ms).

In case of the (coarse-sampled) time series b), binary events need to be extracted from the continuous electrode signal. The extraction of spike times from the continuous signal relies on a criterion to differentiate if the set of observed neurons is spiking or not – which is commonly realized by applying a threshold. (Note that now thresholding takes place on the electrode level, whereas previously, an event belonged to a single neuron.) Here, we obtain avalanches by thresholding as follows: First, all time series are frequency filtered to 0.1 Hz < *f* < 200 Hz. This demeans and smoothes the signal (and reflects common hardware-implemented filters of LFP recordings). Second, the mean and standard deviation of the full time series are computed for each electrode. The mean is virtually zero due to the high-pass filtering. Each electrode’s threshold is set to three standard deviations above the mean. Third, for every positive excursion of the time series (i.e. *V*_*k*_(*t*) > 0), we recorded the timestamp *t* = *t*_max_ of the maximum value of the excursion. An event was defined when *V*_*k*_(*t*_max_) was larger than the threshold *Θk* of three standard deviations of the (electrode-specific) time series. (Whenever the signal passes the threshold, the timestamps of all local maxima become candidates for the event; however, only the one largest maximum *between two crossings of the mean* assigns the final event-time.) Once the continuous signal of each electrode has been mapped to binary events with timestamps, the remaining analysis steps were the same for coarse-sampled and sub-sampled data.

**Table 3:** Fitted exponents of *α* ∼ *Δt*^−*β*^.

| Dynamic state | $\beta$ | |
| --- | --- | --- |
| | $d_E = 200 \mu\text{m}$ | $d_E = 400 \mu\text{m}$ |
| in vitro (LFP) [1] | $0.16 \pm 0.01$ | |
| Critical (coarse) | $0.113 \pm 0.001$ | $0.141 \pm 0.001$ |
| Reverberating (coarse) | $0.127 \pm 0.003$ | $0.156 \pm 0.002$ |
| Subcritical (coarse) | $0.159 \pm 0.004$ | $0.231 \pm 0.016$ |
| Critical (spikes) | $0.143 \pm 0.010$ | $0.123 \pm 0.005$ |

### 4.8 Power-law fitting and shape collapse

Avalanche size and duration distributions are fitted to power-laws using the powerlaw package [58]. The shape collapse of Eq. 5 is done following the algorithm described in [59]. Briefly, the avalanche profiles *s*(*t, D*) of all avalanches with the same duration *D* are averaged, and the resulting curve is scaled to *t*/*D* and interpolated on 1000 points in the [0, 1] range. Avalanches with *D* < 4, or with less than 20 realizations are removed. The chosen collapse exponent *γ* is the one that minimizes the error function:

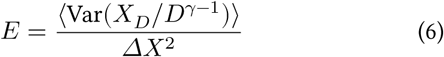

where *X*_*D*_(*t*/*D*) is the interpolated average shape of avalanches with size *D*, and *ΔX* = max_*t,D*_(*X*_*D*_/*D*^*γ*−1^) – min_*t,D*_(*X*_*D*_/*D*^*γ*−1^). The variance Var(.) is calculated over all valid *D*, and the mean ⟨.⟩ is taken over the scaled duration *t*/*D*. For interpolation and minimization we use the scipy [60] functions interpolate.InterpolatedUnivariateSpline and optimize.minimize, respectively.

### 4.9 Data availability

The simulation data used in this study is available from the corresponding author upon request.

## Supplementary Information

### 1.1 Sampling bias remains under alternative topologies

The network topology used in the main paper is local: on average, each neuron is connected to its nearest *K* = 10^3^ neighbors. It is of interest to check if alternative topologies can impact the distinguishability of the underlying dynamic state under coarse-sampling.

For that, we select two additional topologies. The first (“Orlandi”) mimics the growth process of a neuronal culture. In short, axons grow outward on a semiflexible path of limited length and have a given probability to form a synapse when they intersect the (circular) dendritic tree of another neuron. Thereby, this topology is local without requiring distance-dependent synaptic weights (refer to [1] for more details). The second (“Random”) implements a purely random connectivity, with each neuron being connected to *K* = 10^3^ neurons. Note that this is an unrealistic setup as this topology is completely non-local.

We find that, under coarse-sampling, reverberating and critical dynamics remain indistinguishable with the alternative topologies (Fig. S1, left). Meanwhile, under sub-sampling, all dynamic states are clearly distinguishable for all topologies (Fig. S1, right).

### 1.2 Influence of the electrode field-of-view

In the main paper we considered that the contribution of a spiking neuron to the electrode signal decays with distance as ∼ 1/*d*. The precise way neuronal activity is recorded by extracellular electrodes depends on factors such as neuronal morphology and the level of correlation between synapses [2, 3]. Nevertheless, we can study the impact of a varying electrode field-of-view by changing the electrode contribution of a spike to ∼ 1/*dγ* with 1 ≤ *γ* ≤ 2. Note that *γ* = 1 corresponds to an electric monopole, while *γ* = 2 corresponds to an electric dipole – which has a considerably smaller spatial reach.

As *γ* increases, the relative contribution of the closest neurons to the electrode increases, and coarse-sampling becomes more similar to sub-sampling. The cut-off at *S* ∼ *N*_E_ vanishes for large *γ*, and the different dynamic states become distinguishable (Fig. S2D-F). For completeness, in Fig. S4 and Fig. S5 we show the effect of the varying electrode field-of-view for the alternative network topologies discussed previously (“Orlandi” and “Random”), with *d*_E_ = 400 μm and *d*_E_ = 200 μm respectively. In all cases, *γ* ≥ 1.5 results in a vanishing of the cut-off in *p*(*S*). Note, however, that this requires a sufficiently large *d*_E_: for *d*_E_ = 100 μm and *Δt* = 2 ms, an electrode field-of-view of *γ* = 1.5 displays the cut-off, and the dynamic states are not distinguishable (Fig. S2C).

Thus, in order to determine criticality under coarse-sampling, the experimental set-up must combine i) a large *d*_E_, ii) a narrow electrode field-of-view (large *γ*) and iii) systems with different dynamic states. This can potentially then be used to qualitatively compare the distance to criticality between the systems. Not only is this much more limited than what is possible with sub-sampled data [4–6], but the lack of the cut-off is not observed in experimental data of coarse-sampled recordings —which indicate that electrodes typically have a large field-of-view, and that our assumption of *γ* = 0 is adequate.

**Figure S1:**
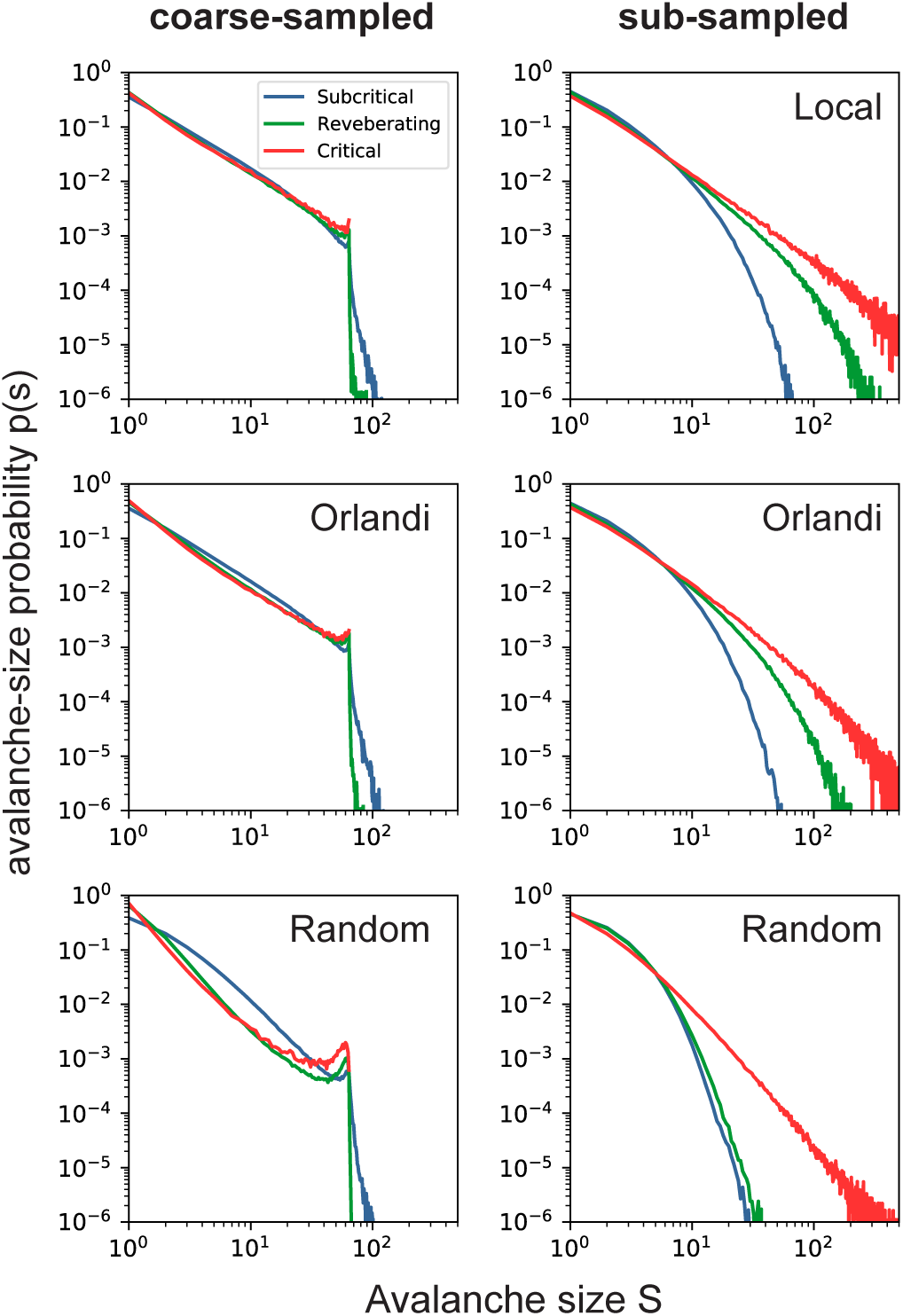
Effect of alternative network topologies. Avalanche-size probability *p*(*S*) from coarse-sampled activity (**left**) and sub-sampled activity (**right**) for subcritical, reverberating and critical dynamics. **Top**: results for the topology used in the main paper (“Local”). **Middle**: results for a topology that mimics culture growth [1] (“Orlandi”). **Bottom**: results for a random topology. Under coarse-sampling, reverberating and critical dynamics are indistinguishable with all topologies. Parameters: *d*_E_ = 400 μm and *Δt* = 8 ms.

### 1.3 Coarse Graining the Ising Model

To demonstrate how general the impact of measurement overlap is, we study the two-dimensional Ising model. The Ising model is well understood and often serves as a text-book example for renormalization group (RG) theory in Statistical Physics [7]. In this framework, the system is *coarse grained* by merging multiple parts (spins) into one. An intuitive way to think of it is by zooming out of a photograph on a computer screen; a pixel can only show one color although there might be more details hidden underneath. Coarse graining is also known as the real-space block-spin renormalization and it can be used to assess criticality. Please note that *coarse graining* is different from *coarse-sampling*. Conventionally, coarse-graining perfectly tiles the space without any measurement-overlap (see Fig. S3).

The two-dimensional Ising model consists of *N* = *L*^2^ spins with states *s*_*i*_ = ±1, arranged on a square lattice of length *L*. In its simplest form, it is given by the Hamiltonian 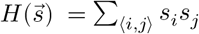, where ⟨*i, j*⟩ denotes all pairs of nearest neighboring spins. The probability of observing 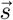 is given by the Boltzmann distribution

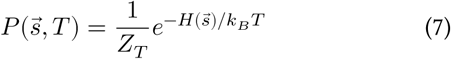

where *T* is the temperature of the system, *k*_B_ is the Boltzmann constant (here, *k*_B_ = 1) and *Z*_*T*_ is the partition function that normalizes the distribution. As the temperature 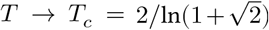, the system undergoes a second-order phase transition between a disordered spin configuration (*T* > *T*_*c*_) and an ordered state of aligned spin orientations (*T* < *T*_*c*_). Many observables diverge at *T* = *T*_*c*_ for *L* → ∞, such as correlation length, specific heat and susceptibility [7, 8].

We perform Monte Carlo simulations of the 2D Ising model using the massively parallel multicanonical method [9, 10]. The multicanonical method offers numerous advantages over conventional Monte Carlo approaches. For instance, instead of simulating at a single temperature, one simulation covers the whole energy space. High-precision canonical expectation values of observables are recovered for any desired temperature during a post-production step. Thereby, we obtain the normalized absolute magnetization as a function of temperature 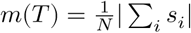.

**Figure S2:**
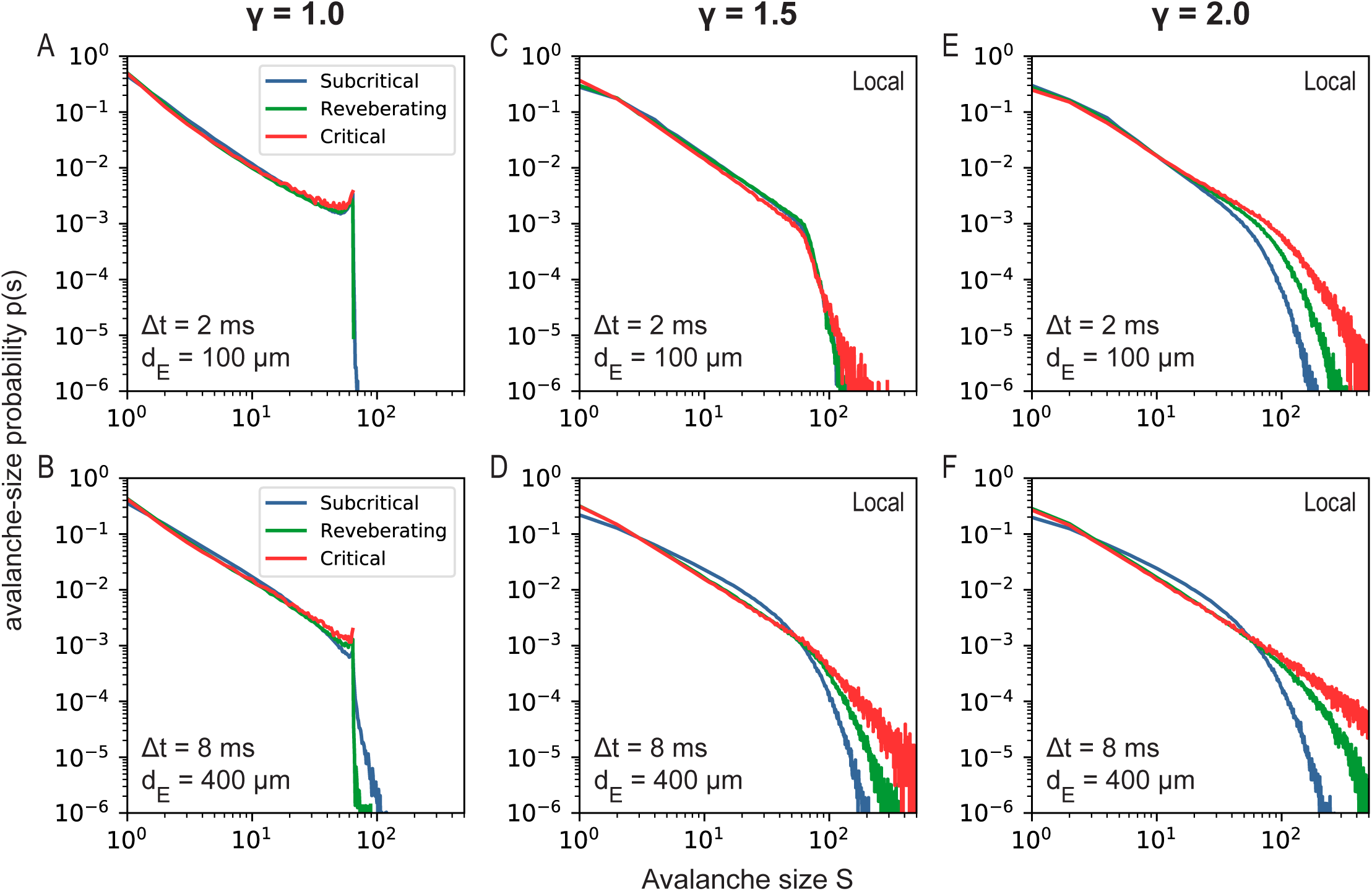
Effect of changing the electrode contribution ∼ 1/*d*^−*γ*^ of a spiking neuron at distance *d*. **A**: Avalanche-size probability *p*(*S*) with *γ* = 1.0 for *Δt* = 2 ms and *d*_E_ = 100 μm. **B**: Avalanche-size probability *p*(*S*) with *γ* = 1.0 for *Δt* = 8 ms and *d*_E_ = 400 μm. **C**: Same as A for *γ* = 1.5. **D**: Same as B for *γ* = 1.5. **E**: Same as A for *γ* = 2.0. **F**: Same as B for *γ* = 2.0. Increasing *γ* results in a smaller electrode field-of-view, and removes the cut-off for *S* ∼ *N*_E_.

### 1.4 Block-Spin Transformation

Measurement overlap causes individual sources to contribute multiple times to a signal. For the Ising model, a similar process takes place when coarse graining is applied. In the process, spins are grouped into blocks of size *b* ×*b*, here *b* = 4 and every block only takes a single value. The value of each block can be obtained in different ways.

- Most commonly, the majority rule [7] is employed, where the block is assigned +1 (−1) if the majority of spins has value +1 (−1). In this case, the contribution of multiple sources is integrated. Hence we compare this rule to the effects observed when neuronal systems are coarse-sampled.
- Alternatively, one can use the decimation rule [7]. In this case, all except a single spin value within a block are discarded. The block value is assigned from the single spin that is kept. Hence we compare this rule to the effects observed when neuronal systems are sub-sampled.

This block-spin transformation rescales the number of spins by a factor of 1/*b*^2^, effectively reducing system size (which will cause finite-size effects). It is well known, that when studying the magnetization, the effective size of the compared systems (after rescaling) has to match.

**Figure S3:**
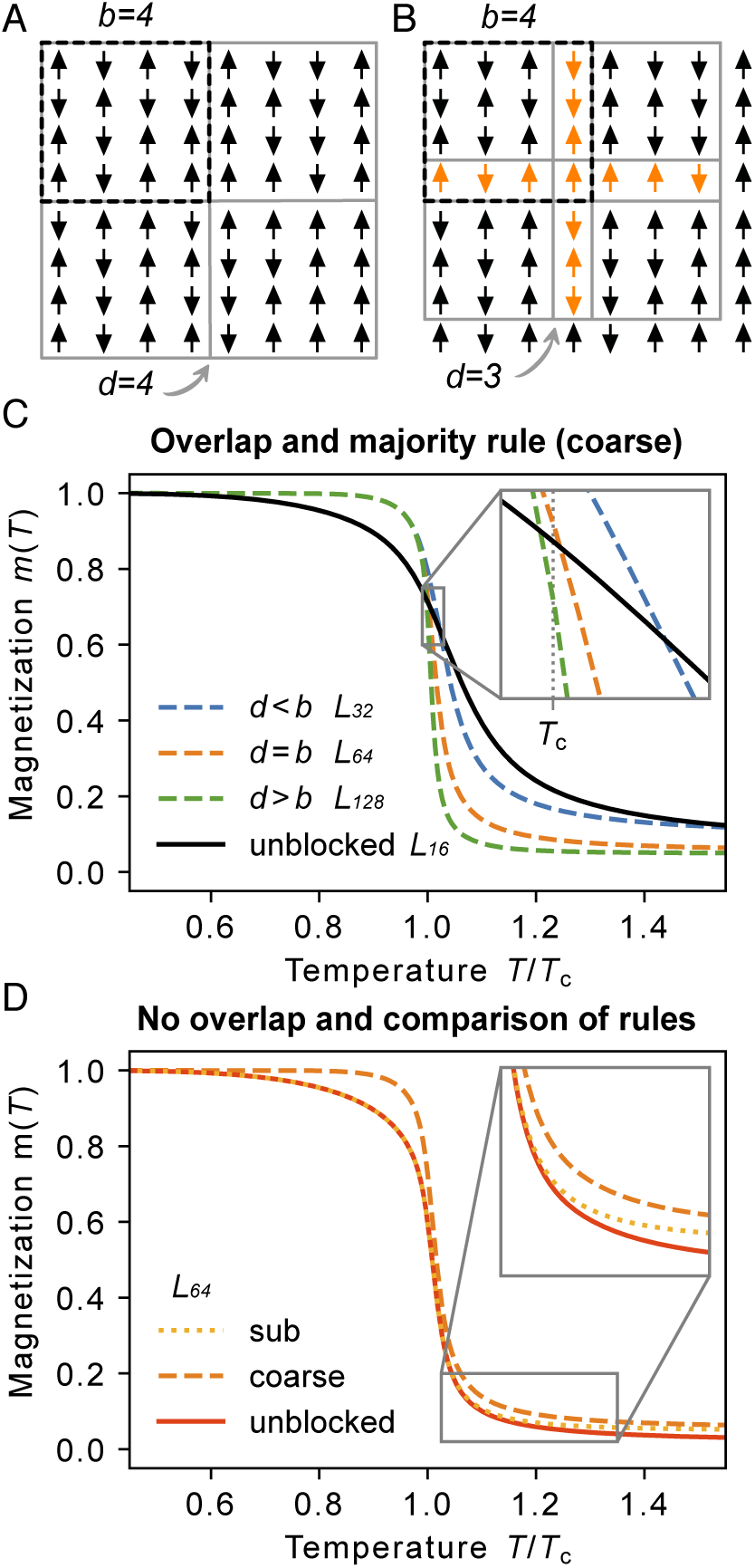
Coarse graining the Ising model. **A:** Representation of the standard coarse graining where block size matches the distance between blocks (*d* = *b* = 4). No overlap is created. **B:** Coarse graining with block size *b* = 4 and a distance between blocks of *d* = 3. Overlapping spins (orange) are shared by two or more blocks. **C:** With the “coarse” majority rule, overlap impacts the spontaneous magnetization *m*(*T*). Only the crossing between the unblocked (*L* = 16) and non-overlapping blocked system (*d* = *b, L* = 64) happens at *T* = *T*_*C*_, as would be expected. Intriguingly, the overlap (*d* < *b, L* = 32) pushes the system towards higher magnetization where spins appear more aligned. On the other hand, the absence of overlap (*d* > *b, L* = 128) causes smaller magnetization where spins appear more random. (Note that, in order to avoid finite-size effects, the target size after coarse graining has to match, here *L* = 16. Consequently, depending on the ratio between *d* and *b*, simulations have different system sizes.) **D:** Comparison between the fully-sampled, unblocked system and blocked systems using the majority rule (“coarse”) and the decimation rule (“sub”) for *d* = *b* = 4. All simulations and curves for *L* = 64. In the ordered, low-temperature phase, the sub curve matches the fully sampled system. Only for the high-temperature phase deviations occur due to finite-size effects (the magnetization for *T* → ∞ approaches the value expected for the rescaled *L* = 16 system). The coarse curve is systematically biased towards more ordered states.

### 1.5 Overlap

To mimic the measurement overlap, we now introduce an overlap between the blocks of the Ising model coarse graining (Fig. S3). In the native block-spin transformation, blocks do not overlap. Then, in terms of spins, the linear distance *d* between two blocks matches the block size *b* = *d* = 4 (Fig. S3A). When the distance between blocks is smaller than the block size, *d* < *b* (Fig. S3B), measurement overlap is created, while when *d* > *b* parts of the system are not sampled. Clearly, the changes that such an overlap will cause on rescaled observables should depend on the rule used perform the block-spin transformation.

Here, we look at combinations of block size *b* = 4 with distance between blocks of *d* = 2, *d* = 4 and *d* = 8. In order to preserve the effective system size (*L* = 16), we thus perform simulations for *L* = 32, *L* = 64 and *L* = 128, respectively.

Using the majority rule and no overlap – which is the default real-space renormalization-group approach – the procedure moves *m* away from *m* (*T*_*c*_) (Fig. S3C, *d* = *b*): For *T* < *T*_*c*_, *m* is increased; For *T* > *T*_*c*_, *m* is decreased. Ordinarily, *T*_*c*_ can be obtained by finding the crossing of *m* between an unblocked (*L* = 16) and a blocked (*L* = 64, *b* = 4) system – only at *T*_*c*_ is the measured *m* invariant under block rescaling transformations.

### 1.6 Majority Rule “coarse”

What is the impact of the overlap for the majority rule? For increasing overlap (*d* < *b*), the crossing occurs at *T* > *T*_*c*_ (Fig. S3C). This is because sharing spins increases the correlations between blocks (pairwise and higher-order), making it more likely that the rescaled spins point into the same direction. In other words, it biases the measurement of *m* towards order, increasing our estimated critical temperature.

For absent overlap (*d* > *b*), only every other block is measured. This decorrelates the spins near the borders of each block and, therefore, decreases the correlation between blocks. As a consequence, the spin orientation of the blocked system moves towards disorder, decreasing the measured magnetization *m*.

### 1.7 Decimation Rule “sub”

If instead of the majority rule the decimation rule is used, the blocking procedure does not alter the correlation between spins before and after the transformation (Fig. S3D). As a consequence, the magnetization remains unaltered in general. However, in the disordered phase, we still notice a systematic deviation from the unblocked system (with *L* = 64). This deviation can be fully attributed to finite-size effects: The distribution of realizable magnetizations in the disordered phase follows a Gaussian with mean zero and variance proportional to the (effective) number of spins. Due to the definition of the magnetization with absolute value, the expectation value of the magnetization for *T* → ∞ is determined by the (effective) system size.

As was the case when sub-sampling neuronal systems, the increase in correlation that ultimately leads to biased observables is caused by integrating weighted contributions from various sources. This is not the case when the decimation rule is applied. Note that the impact of different block-transformation rules on *m*(*T*) will not hold for all other canonical observables such as the energy *E*(*T*) [7].

---

**Figure S4:**
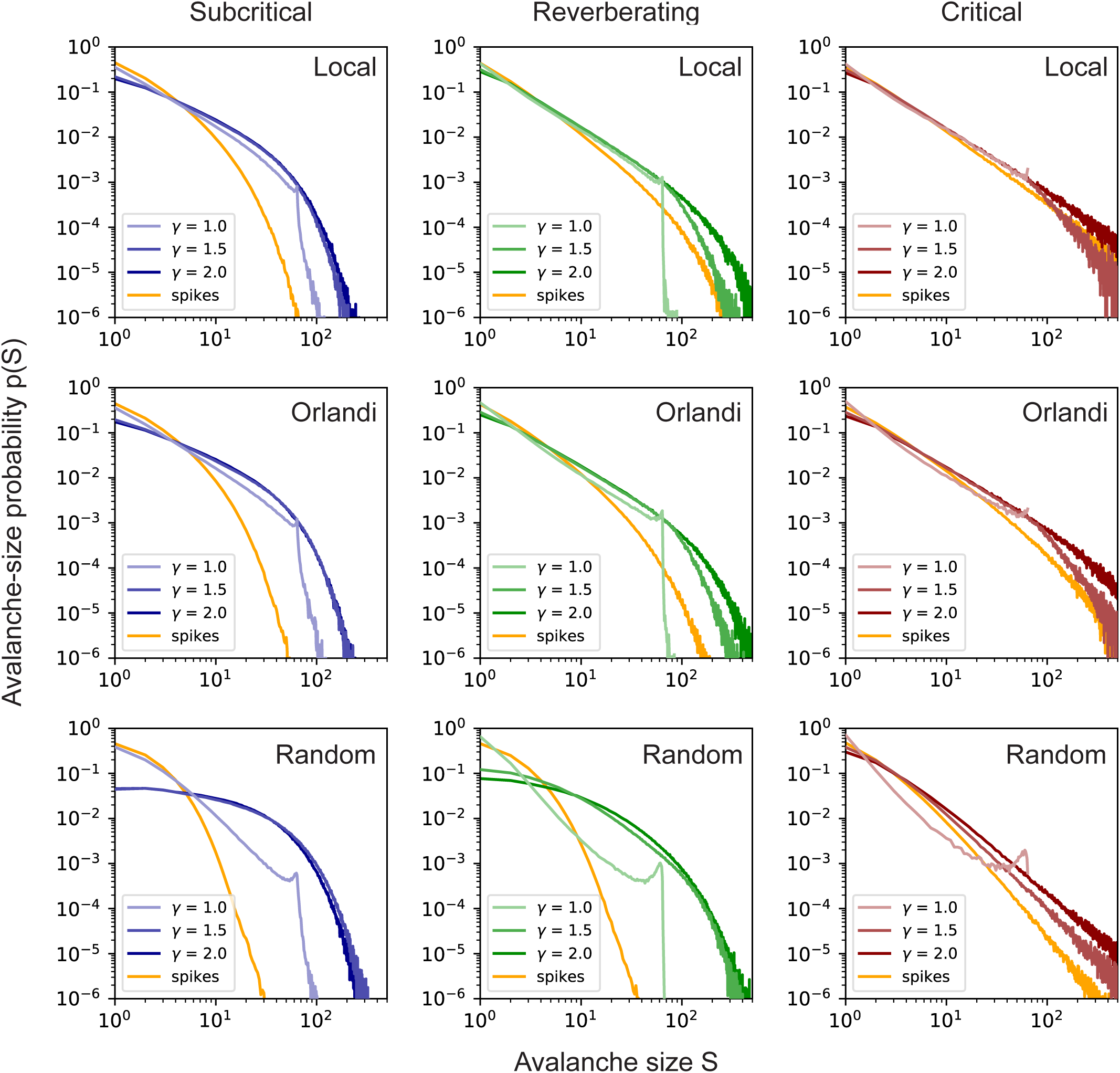
Effect of changing the electrode contribution ∼ 1/*d*^−*γ*^ of a spiking neuron at distance *d*, for different network topologies and *d*_E_ = 200 μm. Dynamic states are Subcritical (**left**), Reverberating (**center**) and Critical (**right**). Topologies are Local (**top**), Orlandi (**middle**) and Random (**bottom**). Local corresponds to the topology used in the main paper, Orlandi corresponds to the model described in [1], and Random corresponds to a completely random topology. Increasing *γ* (decreasing electrode FOV) results in a loss of the cut-off for *p*(*S*) ∼ *N*_E_ as the oarse-sampling becomes more spike-like. Bin-size for all distributions is *Δt* = 4 ms.

**Figure S5:**
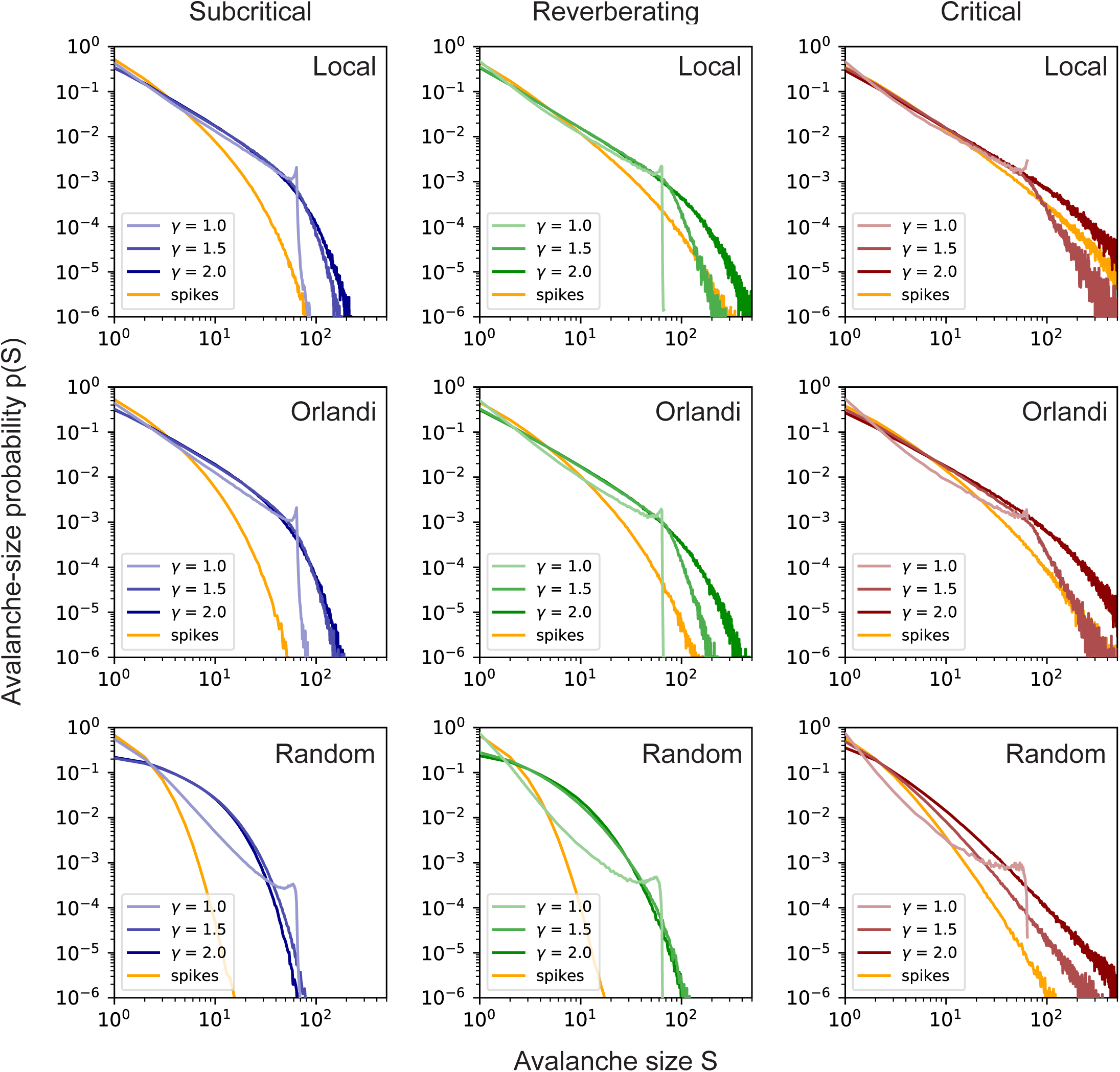
Effect of changing the electrode contribution ∼ 1/*d*^−*γ*^ of a spiking neuron at distance *d*, for different network topologies and *d*_E_ = 400 μm. Dynamic states are Subcritical (**left**), Reverberating (**center**) and Critical (**right**). Topologies are Local (**top**), Orlandi (**middle**) and Random (**bottom**). Local corresponds to the topology used in the main paper, Orlandi corresponds to the model described in [1], and Random corresponds to a completely random topology. Increasing *γ* (decreasing electrode FOV) results in a loss of the cut-off for *p*(*S*) ∼ *N*_E_ as the coarse-sampling becomes more spike-like. Bin-size for all distributions is *Δt* = 8 ms.

**Figure S6:**
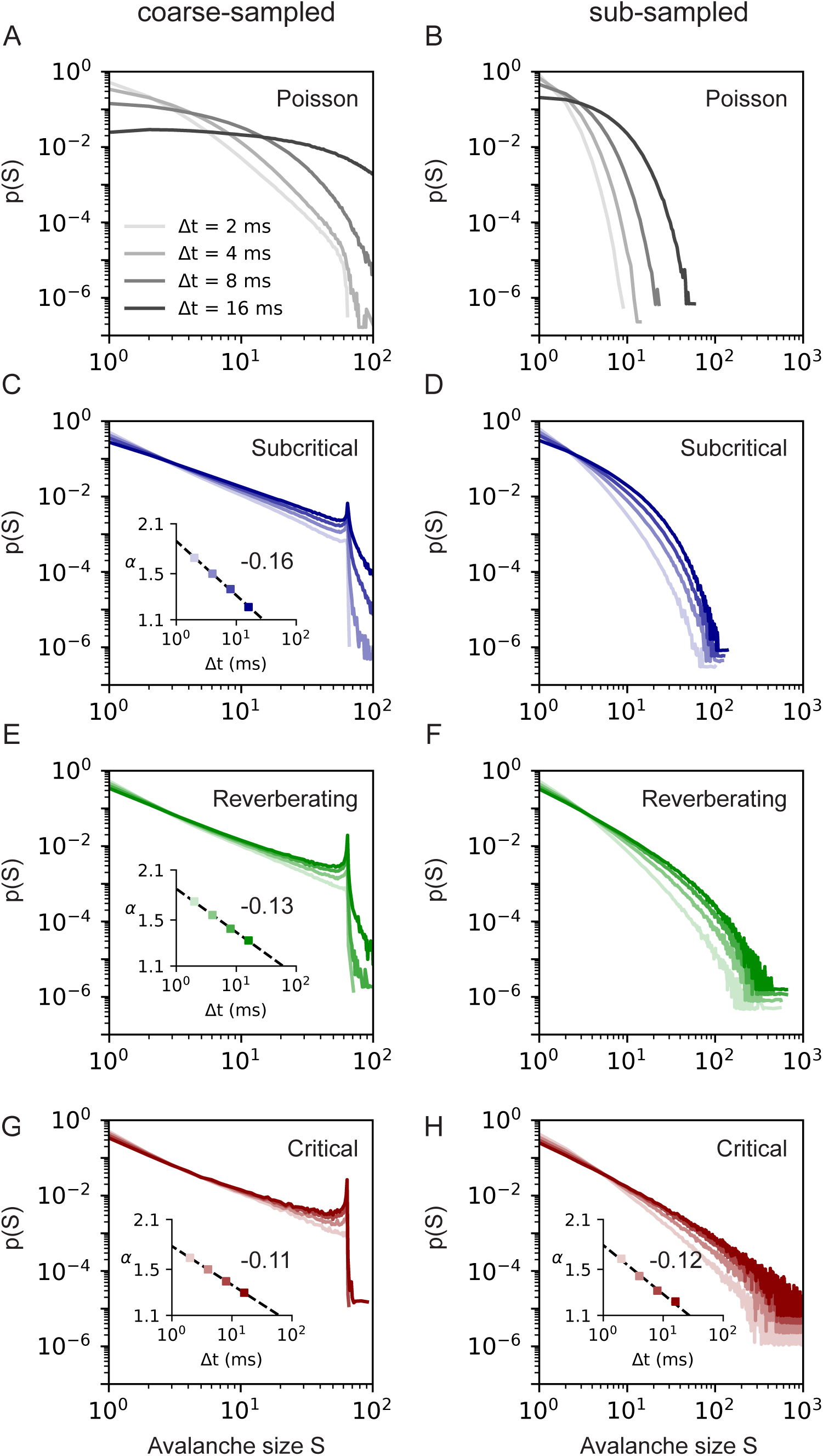
Avalanche-size distributions *p*(*S*) dependence on time-bin size *Δt* for *d*_E_ = 200 μm. Coarse-sampled (**left**) and sub-sampled (**right**) results from an array of 64 virtual electrodes with time bin sizes between 2 ms ≤ *Δt* ≤ 16 ms. Dynamics states are Poisson (**A-B**), Subcritical (**C-D**), Reverberating (**E-F**) and Critical (**G-H**). Distributions are fitted to *p*(*S*) ∼ *S*^−*α*^. **Insets:** Dependence of *α* on *Δt*, fitted as *α* ∼ *Δt*^−*β*^. Fit values are shown in Table. 3.

